# A Bioinformatician, Computer Scientist, and Geneticist lead bioinformatic tool development - which one is better?

**DOI:** 10.1101/2024.08.25.609622

**Authors:** Paul P. Gardner

## Abstract

The development of accurate bioinformatic software tools is crucial for the effective analysis of complex biological data. This study examines the relationship between the academic department affiliations of authors and the accuracy of the bioinformatic tools they develop. By analyzing a corpus of previously benchmarked bioinformatic software tools, we mapped bioinformatic tools to the academic fields of the corresponding authors and evaluated tool accuracy by field. Our results suggest that “Medical Informatics” outperforms all other fields in bioinformatic software accuracy, with a mean proportion of wins in accuracy rankings exceeding the null expectation. In contrast, tools developed by authors affiliated with “Bioinformatics” and “Engineering” fields tend to be less accurate. However, after correcting for multiple testing, no result is statistically significant (*p >* 0.05). Our findings reveal no strong association between academic field and bioinformatic software accuracy. These findings suggest that the development of interdisciplinary software applications can be effectively undertaken by any department with sufficient resources and training.

## Background

Departmental divisions in academia signify research expertise, influence hiring decisions, and impact access to funding, publishing opportunities, and student training [1, 2]. However, interdisciplinary fields such as bioinformatics blur traditional boundaries by integrating data and methods from biology, computer science, and mathematics to address complex research challenges [3, 4, 5].

Bioinformatics has become essential to modern biological research, facilitating evolutionary, structural, and functional analyses of genomic, transcriptomic, and proteomic data [6, 7]. The development of accurate and scalable bioinformatic tools is critical for interpreting such large datasets. Which requires both biological insight and advanced computational skills to build algorithms. The advent of high-throughput technologies has driven the growth of bioinformatics, leading to the establishment of specialized groups within biology, computer science, and engineering faculties, each contributing to the field’s expansion [5, 7].

The contributions of “domain experts” to bioinformatics from the biological and health sciences, such as genetics and molecular biology are essential as they ensure software tools are relevant and accurate. However, domain experts may lack the advanced computational expertise needed to develop sophisticated software. In contrast, fields like mathematics, engineering, and computational sciences — referred to here as “development experts” — offer expertise in algorithm development, mathematical modeling, statistics, and software engineering, essential for creating efficient and scalable bioinformatic tools.

It is possible that departmental differences may influence bioinformatic tool development by reflecting the distinct expertise, resources, and perspectives offered by different academic fields. Development experts excel in computational efficiency, while domain experts provide essential biological insights. Therefore, the success of a tool may depend on the integration of diverse skills rather than on the specific departmental affiliation of its developers.

The accuracy of bioinformatic software tools are of critical importance to the field [8]. Yet several factors can influence accuracy assessments. Previous work has shown that author conflicts can bias tool accuracy assessments upwards [9]. Conversely, commonly assumed features of accurate software such as author reputation (e.g. H-index), tool citation rates, journal impact, runtime, and age of tools had no significant association with software accuracy [10]. Instead, features illustrating sustained development of tools, such as the number of releases, and activity on a popular distributed version control website was significantly associated with accuracy [10].

The primary objective of this study is to examine whether the academic department affiliation of a corresponding author has a discernible outcome on the accuracy (i.e. correctness of predictions) of the bioinformatic tools they develop. Specifically, we aim to determine whether tools created by authors from domain-expert, development-expert of interdisciplinary fields differ in accuracy. This question was motivated by the possibility that certain grant reviewers and other research assessors may base their perception of expertise on departmental affiliation rather than more pertinent information such as research experience. To address this challenge we have analyzed a published corpus of benchmarked bioinformatic tools and evaluated relative accuracy ranks based on the developers’ academic affiliations.

## Results

We explored the relationship between the accuracy of bioinformatic software tools and the academic fields of their developers. Using a previously published corpus of benchmarked accuracy rankings [11], we mapped corresponding authors’ addresses to standardized “fields of study” [12] and grouped them into broader categories. Fields of study is a hierarchical classification that includes specific fields (e.g. “Genetics” or “Mathematics”) or general fields (e.g. “Biological Sciences” or “Mathematics and Statistics”). We further classified these into expertise areas of “Development” (e.g. Computer Science), “Domain” (e.g. Biological Sciences) and “Interdisciplinary” (e.g. Bioinformatics or listing both a Biological and Computational Sciences affiliation).

Figure 1 shows the number of tools (set/intersection sizes) for the general and specific fields (for sets sizes ≥10), as well as the intersections between fields due to corresponding authors with multiple affiliations. E.g. 11 tools have been published by authors who list both a Genetics and Computer Science department as their affiliation. Most bioinformatic tools were developed by authors affiliated with Genetics, Bioinformatics, Computer Science, or similar departments. Among the general fields, Biological Sciences produced the most software tools, followed by Computer Sciences.

**Figure 1.**
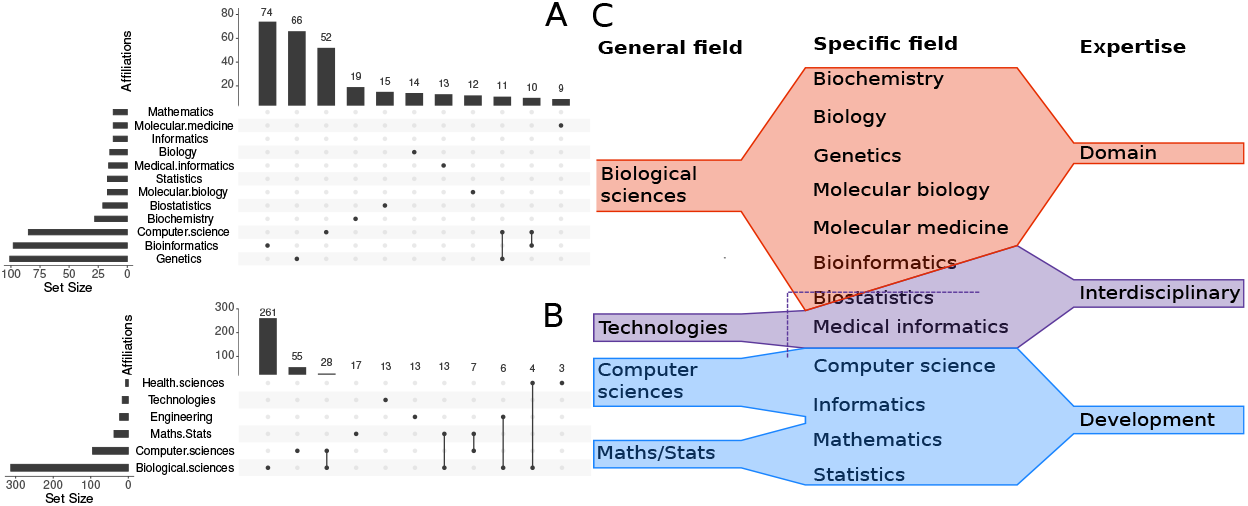
The UpSet figures display the number and intersection of bioinformatic tools classified by general fields (Panel **A**) and specific fields (Panel **B**). Tools are mapped to the affiliations of corresponding authors who published each tool included in selected benchmark studies. Only the top 11 sets are shown for general and specific fields. Panel **C** presents the classification of general and specific fields of study used by the “National Science Foundation” relevant to this study. On the right, it shows the additional expertise categories that we introduced for this analysis.

We ranked fields based on the mean proportion of “wins” (i.e., for field “1” when its tool “A” outperforms tool “B” in benchmark “X”) and calculated Z-scores to compare them against random expectations (i.e., *wins* = 0.5). A higher proportion of wins and lower (−1) ∗*Z* − *score* indicate better overall accuracy.

“Medical Informatics”, a branch of “Technologies”, out-performed other fields, with a mean win proportion of 0.70 (95% CI: 0.53 − 0.85) and a Z-score of −1.88. However, P = 0.29 after multiple testing correction. Notably, this category includes five different parameter options for the MAFFT sequence alignment tool [13], and a further four separate corresponding authors that list either the Department or Center of Biomedical Informatics, Harvard Medical School as their affiliation [14, 15, 16, 17]. This and some other redundancies leaves just eight departments representing the medical informatics field here.

In contrast, “Bioinformatics” has the lowest rank, with a mean win proportion of 0.43 (95% CI: 0.33 − 0.53) and a Z-score of 1.20 (P = 0.46 after correction). Similarly, “Engineering” ranked low, with a win proportion of 0.34 (95% CI: 0.16 − 0.60) and a Z-score of 1.25.

Other fields showed confidence intervals that included the null value of 0.5, with modest Z-scores ranging from −0.49 to 0.96, and P-values greater than 0.05.

When grouped by expertise type—software development experts, biological domain experts, or interdisciplinary experts—all categories had similar win proportions (0.51, 0.49, and 0.46, respectively). Interdisciplinary experts had the lowest Z-score of −0.87 (P = 0.46 after correction).

## Conclusions and Limitations

We tested the assumption that academic department specialization reflects the quality of research software. After correcting for multiple testing, we found no significant association between academic expertise and the accuracy of bioinformatic tools. This suggests that department affiliation does not correlate with software quality, and neither general nor specific research fields showed any significant links to tool accuracy.

A previous study indicated that maintaining software was the key factor in producing accurate tools, while citation metrics, tool age, speed are not associated with software accuracy [10]. Our findings complement this by showing that academic field is also not associated with software accuracy.

While other aspects of bioinformatic tools, such as speed and usability, are important, we emphasize that accuracy should remain the top priority, as poor predictions have long-term impacts on research [8].

Medical Informatics was the top-performing field in developing accurate tools, these include methods for structural variation detection, single-cell profiling, long-read assembly, multiple sequence alignment and are derived from a limited number of research teams. However, tools from Bioinformatics and Engineering ranked lower, though these differences were not statistically significant.

Therefore, an individual’s department is not a reliable indicator of the quality of the software they produce. Academic affiliation should not be used as a proxy for assessing the potential success of software development projects.

### Limitations

Some benchmarks include multiple tool options, potentially introducing non-independent effects. The accuracy metrics are diverse, with some limitations (e.g., issues with “accuracy” in class-imbalanced datasets [18] and criticisms of the N50 metric for sequence assembly [19]). Additionally, smaller benchmarks and smaller fields may exaggerate rank shifts. We mitigate this in part by only considering departments with 10 or more corresponding tools.

Potential confounding variables may exist for this analysis. However, a prior analysis of this data [10] found no correlation between software accuracy and factors such as citations for tools, authors or journal, as well as tool speed, or age, allowing these variables to be ruled out as confounders.

There may be little connection between a researcher’s training and their listed department, as illustrated by this author’s background which began in mathematics, before taking positions in departments of computer science, molecular biology and bioinformatics, and now is affiliated with a Biochemistry Department.

The corresponding (last) author, is typically the principal investigator and may not be the primary tool developer, but rather oversees the project. There is likely to be a strong overlap between the departments of the first and last authors, but this was not explored in the current study.

### Final words

This study does not find evidence linking academic department affiliation with bioinformatic software accuracy (*p >* 0.05 in all instances). Future research should investigate other factors, such as interdisciplinary collaborations and developer training, to understand what drives high-quality tool development. Addressing potential biases against interdisciplinary work [20] and ensuring long-term support for essential software infrastructure will also be critical for advancing the field [21].

## Methods

The data, scripts, figures and manuscript draft files are availble at the GitHub repository: https://github.com/ppgardne/departments-software-accuracy

### Pre-registration

The methods for this study followed the pre-registered proposal outlined prior to any unpublished data collection [11].

### Benchmarking data

software ranks from previously gathered benchmarks are publically available [10], these include data from 68 publications, 128 benchmark rankings of different sets of 498 distinct software tools (summarised in Supplementary Table 1).

In brief, the criteria for inclusion of a benchmark study are based on the definition of “neutral comparison studies” [22]: 1. The main focus of the article is the comparison and not the introduction of a new tool. 2. The authors should be neutral (i.e. not tool developers). 3. The test data and evaluation criteria should be sensible.

### Mapping tools to academic field

For each software tool, the corresponding publication(s) were identified, and the addresses of the primary corresponding author were manually extracted when available. If an author listed multiple addresses, only the first two were used. In cases with multiple corresponding authors, the last corresponding author was chosen.

The department names of the authors were mapped to the closest associated “fields of study” as defined by the National Science Foundation [12]. We analysed these fields at three hierarchical levels: first, specific fields (e.g. “genetics”, “computer science”, “bioinformatics” etc), which were then mapped to broader general fields (e.g. “biological sciences”, “computer sciences” etc). Thirdly, we categorized them into three types of expertise: **development experts, domain experts and interdisciplinary experts**. Development experts, from fields such as computer science, mathematics, and engineering, are expected to bring relevant expertise in software engineering and the mathematical modeling of biological problems. Domain experts, from the biological and health sciences, are anticipated to possess detailed knowledge of their subject area and to be invested in producing high-performing software for their research needs. Interdisciplinary experts come from fields such as bioinformatics, biostatistics, and biomathematics, and also include researchers who list both development and domain expertise (e.g. “Computer Science” and “Genetics”). We have treated some fields as synonymous; for example, “Computational Biology” was mapped to “Bioinformatics”, and “Genomics” is mapped to “Genetics”.

We restricted all subsequent analyses to fields that contain at least 10 software tools in our benchmark corpus. This mitigated against potential issues due to small sample sizes.

### Statistical analysis

The accuracy data is derived from benchmarks using a diverse number of metrics that include sensitivity, specificity, PPV, FDR, error rates, AUROC, MCC and others [8]. The number of tools ranked in any benchmark ranged from 3 to 50. In order to obtain a representative measure of accuracy for a field that accounts for the diversity in accuracy measures and number of ranked tools, we employed a rank-based and bootstrapping strategy. We randomly sampled, with replacement, sets of 200 tools from the total of 498 tools. For each tool, a corresponding benchmark was selected at random, and the number of times the tool “won” against another tool was recorded, along with the total number of pairwise comparisons made. These counts of wins and total comparisons were then assigned to the corresponding specific and general departments, and expertise areas. In other words, a tool ranked second in a benchmark of 11 tools will contribute 9 wins and 10 comparisons to the totals for its corresponding fields.

This process was repeated 1,000 times to estimate the mean proportions of wins for each field, along with a 95% confidence interval for these values (Figure 2A). Additionally, we calculated a Z-score for each field to determine the number of standard deviations the mean number of wins deviates from the expected null value of 0.5 for randomly grouped tools (Figure 2B).

**Figure 2.**
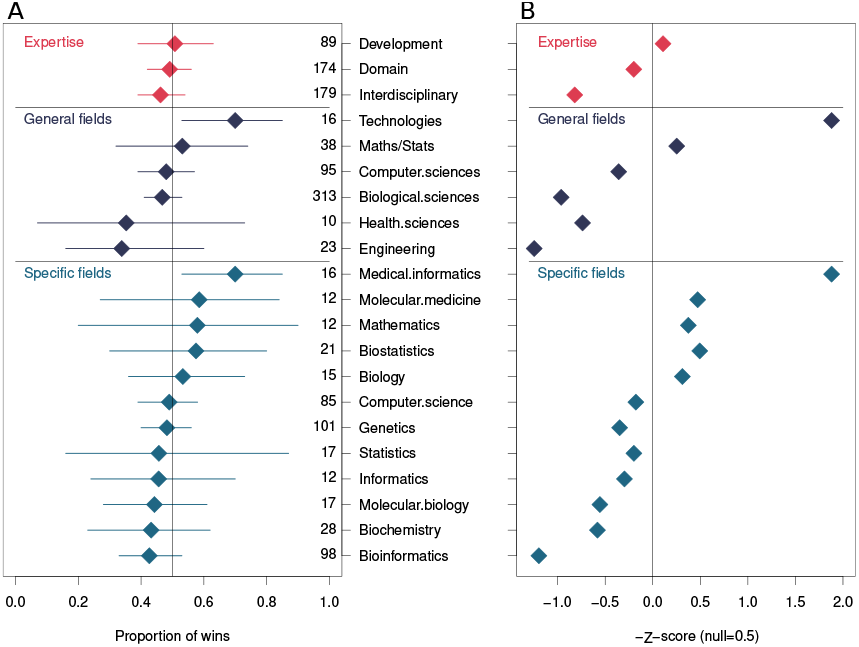
(**A**) A forest plot, illustrating the mean and 95% confidence intervals of the proportion of times software tools published by a given field “win” in pairwise comparisons. Confidence intervals and the mean was determined using a bootstrapping procedure. Within each field the entries have been sorted by the mean number of wins. The sample size for each field is indicated by the column of numbers on the right of the figure. (**B**) A Z-score was computed for each distribution of bootstrap samples for each field. The expected proportion of wins for randomly selected groups of tools was used as “*x*” (i.e. null=0.5).

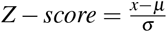

Where *µ* is the mean, *σ* is the standard deviation, *x* is the raw value. In this case we set *x* = 0.5 as this is the null expectation for the proportion of wins for randomly grouped sets of tools. For the purposes of illustration we plot (−1) ∗*z* so that the direction is the same as for the “proportion of wins” forest plot (Figure 1).

P-values are computed from the absolute value of the Z-scores to evaluate if any field is significantly distinguished from the null i.e. *P*[*X > x*]. The P-values are corrected for multiple testing by controlling the false discovery rate method [23].

## Acknowledgements

This research was supported by the MBIE data science platform “Beyond Prediction: explanatory and transparent data science”, Genomics Aotearoa, and Marsden Grants 19-UOO-040 and 20-UOA-282.

Associate Professor Anthony Gitter (Department of Bio-statistics and Medical Informatics, University of Wisconsin-Madison) for pointing out the over-representation of Harvard Medical School affiliations in the Medical Informatics catagory.

The author thanks the anonymous grant reviewers who inspired this paper.

## References

[1] Paul Bourke and Linda Butler. Institutions and the map of science: matching university departments and fields of research. Research Policy, 26(6):711–718, 1998.

[2] Ben R. Martin. What’s happening to our universities? Prometheus, 34:7–24, 2016.

[3] C A Ouzounis and A Valencia. Early bioinformatics: the birth of a discipline–a personal view. Bioinformatics, 19(17):2176–90, Nov 2003.

[4] S R Eddy. “antedisciplinary” science. PLoS Comput Biol, 1(1):e6, Jun 2005.

[5] Paulien Hogeweg. The roots of bioinformatics in theoretical biology. PLoS computational biology, 7(3):e1002021, 2011.

[6] Levin Clement, Dynomant Emeric, Mouchard Laurent, Landsman David, Hovig Eivind, Vlahovicek Kristian, et al. A data-supported history of bioinformatics tools. arXiv preprint 1807.06808, 2018.

[7] Jeff Gauthier, Antony T Vincent, Steve J Charette, and Nicolas Derome. A brief history of bioinformatics. Briefings in bioinformatics, 20(6):1981–1996, 2019.

[8] Lukas M Weber, Wouter Saelens, Robrecht Cannoodt, Charlotte Soneson, Alexander Hapfelmeier, Paul P Gardner, Anne-Laure Boulesteix, Yvan Saeys, and Mark D Robinson. Essential guidelines for computational method benchmarking. Genome biology, 20:1–12, 2019.

[9] Stefan Buchka, Alexander Hapfelmeier, Paul P Gardner, Rory Wilson, and Anne-Laure Boulesteix. On the optimistic performance evaluation of newly introduced bioinformatic methods. Genome biology, 22(1):152, 2021.

[10] P P Gardner, J M Paterson, S McGimpsey, F Ashari-Ghomi, S U Umu, A Pawlik, A Gavryushkin, and M A Black. Sustained software development, not number of citations or journal choice, is indicative of accurate bioinformatic software. Genome Biol, 23(1):56, Feb 2022.

[11] P P Gardner. Pre-registration: Which department makes the best software?, 2024. 10.17605/OSF.IO/92PTZ [10 July 2024].

[12] IPEDS Completions Survey; National Center for Science Department of Education, National Center for Education Statistics and Survey of Earned Doctorates. Engineering Statistics. Classification of fields of study, 2014. https://ncsesdata.nsf.gov/sere/2018/html/sere18-dt-taba001.html [Accessed: July 2024].

[13] Kazutaka Katoh and Hiroyuki Toh. Recent developments in the MAFFT multiple sequence alignment program. Briefings in bioinformatics, 9(4):286–298, 2008.

[14] Tae-Min Kim, Lovelace J Luquette, Ruibin Xi, and Peter J Park. rSW-seq: algorithm for detection of copy number alterations in deep sequencing data. BMC bioinformatics, 11:1–13, 2010.

[15] Lixing Yang, Lovelace J Luquette, Nils Gehlenborg, Ruibin Xi, Psalm S Haseley, Chih-Heng Hsieh, Chengsheng Zhang, Xiaojia Ren, Alexei Protopopov, Lynda Chin, et al. Diverse mechanisms of somatic structural variations in human cancer genomes. Cell, 153(4):919–929, 2013.

[16] Peter V Kharchenko, Lev Silberstein, and David T Scadden. Bayesian approach to single-cell differential expression analysis. Nature methods, 11(7):740–742, 2014.

[17] Jue Ruan and Heng Li. Fast and accurate long-read assembly with wtdbg2. Nature methods, 17(2):155–158, 2020.

[18] Amalia Luque, Alejandro Carrasco, Alejandro Martín, and Ana de Las Heras. The impact of class imbalance in classification performance metrics based on the binary confusion matrix. Pattern Recognition, 91:216–231, 2019.

[19] Luyu Xie and Limsoon Wong. PDR: a new genome assembly evaluation metric based on genetics concerns. Bioinformatics, 37(3):289–295, 2021.

[20] Lindell Bromham, Russell Dinnage, and Xia Hua. Interdisciplinary research has consistently lower funding success. Nature, 534(7609):684–687, 2016.

[21] A Siepel. Challenges in funding and developing genomic software: roots and remedies. Genome biology, 20(1):1–14, 2019.

[22] Anne-Laure Boulesteix, Sabine Lauer, and Manuel JA Eugster. A plea for neutral comparison studies in computational sciences. PloS one, 8(4):e61562, 2013.

[23] Yoav Benjamini and Yosef Hochberg. Controlling the false discovery rate: a practical and powerful approach to multiple testing. Journal of the Royal Statistical Society: Series B (Methodological), 57(1):289–300, 1995.

